# Second-order Citations in Altmetrics: A Case Study Analyzing the Audiences of COVID-19 Research in the News and on Social Media

**DOI:** 10.1101/2023.04.05.535734

**Authors:** Juan Pablo Alperin, Alice Fleerackers, Michelle Riedlinger, Stefanie Haustein

## Abstract

The potential to capture the societal impact of research has been a driving motivation for the use and development of altmetrics. Yet, to date, altmetrics have largely failed to deliver on this potential because the primary audience who cites research on social media has been shown to be academics themselves. In response, our study investigates an extension of traditional altmetric approaches that goes beyond capturing direct mentions of research on social media. Using research articles from the first months of the COVID-19 pandemic as a case study, we demonstrate the value of measuring ‘second-order citations,’ or social media mentions of news coverage of research. We find that a sample of these citations, published by just five media outlets, were shared and engaged with on social media twice as much as the research articles themselves. Moreover, first-order and second-order citations circulated among Twitter accounts and Facebook accounts that were largely distinct from each other. The differences in audiences and engagement patterns found in this case study highlight the importance of news coverage as a public source of science information and provide strong evidence that investigating these second-order citations can be an effective way of observing non-academic audiences that engage with research content.

## Introduction

Since their inception, altmetrics, in particular those based on social media (Sugimoto et al., 2017) were introduced with the hope of identifying the societal impact of research (Bornmann, 2016; Costas et al., 2021; Kassab et al., 2020; Robinson-Garcia et al., 2018). The potential of altmetrics for this purpose lies in how they capture the research being circulated by individuals who might not be writing research papers and who can therefore not be observed through traditional bibliometric analyses (Haustein et al., 2016). Researchers have been especially interested in identifying non-academic audiences who share research on social media platforms that are used by a broad segment of society, such as Twitter and Facebook (Alperin et al., 2019; Costas et al., 2020; Díaz-Faes et al., 2019; Haustein, 2019).

Even if a large number of studies have shown that correlations between social media mentions and citations are low (Costas et al., 2015; Haustein et al., 2014, 2015), there is little indication that these differences stem from different user populations. That is, most studies of social media user communities find that the activity around research articles on social media is largely generated by academics, not members of the general public (Alperin et al., 2019; Carlson & Harris, 2020; Toupin & Haustein, 2018; Tsou et al., 2015; Vainio & Holmberg, 2017). Ferreira, Mongeon & Costas (2021) found that researchers tend to tweet about similar subjects they publish on, indicating that their Twitter activity is an extension of their scholarly communication (Costas et al., 2021). Therefore, despite the hope that social media metrics would be able to capture societal impact of research, they seem to reflect online engagement by academic users.

As such, altmetrics in their current form have not been able to deliver on their promise to capture the broader impact of research in society, or even provide an understanding of what research circulates in the public sphere, and among whom. Given the apparent low volume of research shared on social media by non-academic audiences (Carlson & Harris, 2020), it is necessary to look beyond the mentions of research outputs themselves and to look to other places where research circulates among non-academic audiences. One such place is the news media, which plays a critical role in shaping public discourse (Gallagher et al., 2021; McCombs, 2002) and remains a key source of public science information (Covens et al., 2018; Funk et al., 2017). Metrics that capture news coverage of research may offer insights into its societal impact that have not been captured through other altmetrics (Casino, 2018; Fleerackers, Nehring, et al., 2022; Ortega, 2020a).

Although companies such as Altmetric and PlumX now enable researchers and publishers to calculate and track mentions of research in the news media, only a few studies have sought to use these metrics for understanding the audiences of research (e.g., Maggio et al., 2019; Matthias et al., under review; Moorhead et al., 2021). These few studies have yielded some insights into the diverse media outlets that report on research and the journalistic approaches they use to do so. As such, they serve as a good starting point for understanding the role of the media in mobilizing research knowledge to a broad, nonacademic public. However, they only examine journalistic attention to the original journal articles, and not the wider attention given to those news stories within society.

One way to address this shortcoming is by exploring so-called “second-order citations,” or social media posts that mention (i.e., cite) web pages that cite research (e.g., news stories that mention research articles) (Priem & Costello, 2010). Unlike typical altmetrics that focus on “first-order citations” (i.e., social media posts that link to research directly), second-order citations provide the opportunity to observe a common way for non-academic users to share research with friends or followers (Lemke et al., 2021). While the impact of these citations is not yet fully understood, it is evident that news stories mentioning research have the potential to reach users who would not otherwise engage with research on social media, amplifying academic knowledge to broad audiences (Fleerackers, Riedlinger, et al., 2022). This study seeks to investigate the impact of these second-order citations through an analysis of tweets and Facebook posts linking the original research articles as well as news stories about the article. It asks: *How does social media engagement with news stories about research compare to engagement with the research articles themselves?* Building on a unique and novel dataset, this study treats COVID-19-related research as a case study through which to exemplify the methodological approach and point to its potential value.

## Methods

### Research Articles

In March 2021, we identified all COVID-19-related articles published between January 1, 2020 and December 31st, 2020 using a well-established set of search terms from the National Library of Medicine (Chen et al., 2020; see Appendix). We restricted our search to articles from two preprint servers (*bioRxiv* or *medRxiv*), because they were prolific sources of COVID-19-related research during the first year of the pandemic, and two peer-reviewed journals (*Journal of Virology* or *British Medical Journal*), because they were among the most active players publishing COVID-19 research and sped up their peer review process in the beginning of the pandemic (Kousha & Thelwall, 2020; Palayew et al., 2020). In total, we identified 3,934 relevant research articles within these four outlets.

### News Stories

We collected news stories that mentioned any of the research articles in our sample by querying the Altmetric Explorer. Altmetric is a company that collects mentions of scholarly documents in online news and social media by regularly scanning the text of thousands of media stories. It identifies news stories that mention research articles either through links to the publication (i.e., through a URL or a publication identifier, such as a DOI) or mentions of study details such as author names, journal titles, and publication dates (Altmetric.com, 2018). While these data are not without issues (Ortega, 2019, 2020b), they can be reasonably accurate when working with a fixed set of news outlets (Fleerackers, Nehring, et al., 2022). With this limitation in mind, we restricted our query to five outlets that circulate widely on Twitter and Facebook: *BBC, MSN, The New York Times, The Guardian*, and *The Washington Post*. This search revealed that 344 (8.7%) of the research articles in our sample were mentioned 1,406 times across 1,221 unique news stories (a news story can mention several different research outputs; see Table 1). On average, each article was mentioned 4.1 times (SD = 6.5).

**Table 1.** Number of news stories citing research articles in five outlets

|  | Number of Stories Citing Research |  |  |  |  | Total | Number of Research Articles |
| --- | --- | --- | --- | --- | --- | --- | --- |
|  | BBC | MSN | The New York Times | The Guardian | The Washington Post |  |  |
| **British Medical Journal** | 47 | 262 | 33 | 73 | 22 | 437 | 147 |
| **Journal of Medical Virology** | 11 | 85 | 43 | 4 | 7 | 150 | 39 |
| **bioRxiv** | 5 | 116 | 92 | 10 | 10 | 233 | 43 |
| **medRxiv** | 20 | 357 | 143 | 36 | 30 | 586 | 115 |
| **Number of stories** | 83 | 820 | 311 | 123 | 69 | 1406 | 344 |

### Social Media Mentions

To search for social media posts with links to the research articles (i.e., first-order citations), we identified three possible URL types where each research article might be found. The first two URLs were based on Crossref’s guidelines for creating links from an article’s Digital Object Identifier (DOI), using the patterns http://dx.doi.org/{doi} and https://doi.org/{doi} (Hendricks, 2017). A third URL was identified by resolving the DOI URL using a Python script. Similarly, we resolved the shortened URLs provided by Altmetric to arrive at the URL to each of the news stories. To identify social media posts with news stories that mentioned those articles (i.e., second-order citations), we used the URLs provided by Altmetric.

Between March 9 and April 9, 2021, we used Python scripts, including the Twint library (Poldi & Zacharias, 2017/2021), to collect all tweets posted between January 1, 2020 and January 31, 2021 that contained a link to any of the 344 research articles mentioned in the five news outlets or with a link to any of the 1,221 news stories that mentioned those articles. This search yielded 50,299 tweets linking to 325 (94.5%) of the research articles and 97,235 tweets mentioning 486 (39.8%) of the news stories. During the same time period, we used the social media analytics tool Crowdtangle to extract publicly accessible Facebook posts from profiles, groups, and pages ((i.e., public Facebook “spaces”; Bruns et al., 2020) that contained links to the research articles or news stories. While public spaces represent only a small proportion of Facebook attention to research (Enkhbayar et al., 2019), for ethical reasons, Crowdtangle only provides data related to Facebook community activity. This search yielded 6,420 Facebook posts linking to 246 (71.5%) of the research articles and 14,081 posts linking to 516 (42.3%) of the news stories.

The final dataset comprised four elements: 1) 344 research articles; 2) 1,221 news stories that mentioned those research articles; 3) 50,299 tweets and 6,420 Facebook posts that linked to the research articles (i.e., first-order citations), and 4) 97,235 tweets and 14,081 Facebook posts that linked to the news stories (i.e., second-order citations) (Figure 1). Some news stories and social media posts cited more than one research article in our sample, and some social media posts cited more than one news story.

**Figure 1.**
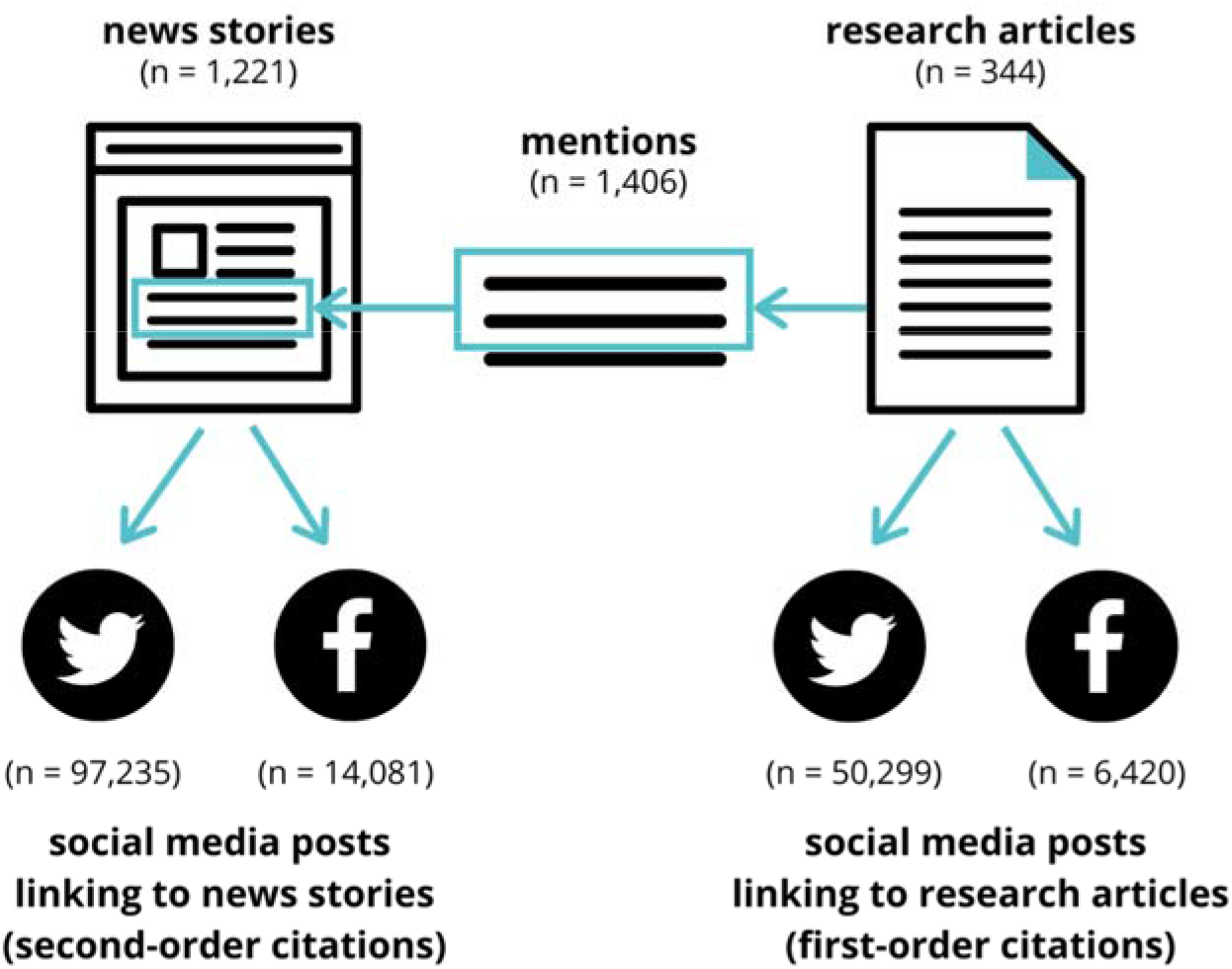
Description of dataset and relationships between each component

### Comparison of Twitter users with known researchers on Twitter

In an effort to estimate whether or not the share of researchers was higher among Twitter user accounts tweeting research articles in comparison to those users who shared the news stories, we determined the overlap between user accounts in our dataset and those identified as researchers in a dataset published by Mongeon, Bowman & Costas (2022, 2023). This dataset contains over 400,000 Twitter accounts that are believed to belong to a researcher based on matching names and a record of having linked to one research article with an author sharing the same name. Mongeon et al. (2023)’s approach was limited to identifying accounts that had previously shared their own research, but was demonstrated to have high precision and moderate recall.

### Statistical Methods and Scripts

All statistics and correlations between first-order citations and second-order citations were calculated using the Python Pandas package (The Pandas Development Team, 2023). For each research article, we calculated the sum of Twitter and Facebook posts linking to the article (first-order citations) and compared these totals to the sum of corresponding posts linking to news stories that mentioned that article (second-order citations). For Twitter, we compared the Twitter user IDs of those who shared each research article with those who shared a news story citing that article. Similarly, for Facebook, we compared the account IDs of spaces (i.e., group, page, or profile) where each research article was posted to those where news stories were posted. All Python scripts written to download data, expand URLs, and perform analyses are available online at Alperin (2023b). All data is available at Alperin (2023a)

## Results

### Size of audiences of research and news

Our analysis shows that second-order citations (i.e. news stories reporting on the research) are shared more frequently on Twitter and Facebook than first-order citations of the research. Collectively, the news stories written by the five media outlets analyzed were shared approximately twice as often as the research articles and by approximately twice as many unique accounts. The news stories also received approximately twice as much engagement as measured in retweets/shares, likes/reactions, and replies/comments) (Table 2).

**Table 2.** Social media attention to research and to news stories mentioning the research

| Twitter |  |  | Facebook |  |  |
| --- | --- | --- | --- | --- | --- |
|  | First-order<br>(Research) | Second-order<br>(News) |  | First-order<br>(Research) | Second-order<br>(News) |
| **Tweets** | 50,299 | 97,235 | **Posts** | 6,420 | 14,081 |
| **Accounts** | 27,771 | 62,290 | **Spaces** | 3,976 | 8,191 |
| **Retweets** | 227,041 | 412,509 | **Shares** | 89,422 | 412,104 |
| **Likes** | 512,308 | 1,111,458 | **Reactions** | 176,890 | 1,476,174 |
| **Replies** | 39,788 | 89,509 | **Comments** | 36,203 | 304,614 |

High Spearman correlations between Facebook shares and tweets for news (*ρ* = 0.95) and research articles (*ρ* = 0.84) shows that patterns on both social media platforms were similar (Table 3). However, when comparing first- to second-order citations, the correlations were very low, even when comparing posting about news articles with posting about research articles on the same platform (*ρ* = 0.13 for tweets and *ρ* = 0.02 for Facebook posts). Correlations were even lower when calculated for activity across platforms (*ρ* = −0.01 for first-order Facebook posts and second-order tweets; *ρ* = 0.00 for first-order tweets and second-order Facebook posts). These relationships are visualized in Figure 2. Each scatterplot represents the relationship between the two variables indicated, with a positive correlation clearly visible for research tweets and Facebook posts (first column, second row) and for news tweets and Facebook posts (bottom row, second column).

**Table 3.** Spearman correlations (ρ) between social media posts mentioning the research article (first-order) and the news stories mentioning that research (second-order)

|  | First-order<br>Twitter (Research) | Second-order<br>Twitter (News) | First-order<br>Facebook<br>(Research) | Second-order<br>Facebook (News) |
| --- | --- | --- | --- | --- |
| **First-order Twitter (Research)** |  |  |  |  |
| Second-order Twitter (News) | 0.13 ($p = .0157$) | | | |
| First-order Facebook (Research) | 0.84 ($p < .0001$) | -0.01 ($p = .8720$) | | |
| Second-order Facebook (News) | 0.02 ($p = .7058$) | 0.95 ($p < .0001$) | 0.00 ($p = .9439$) | |

**Figure 2.**
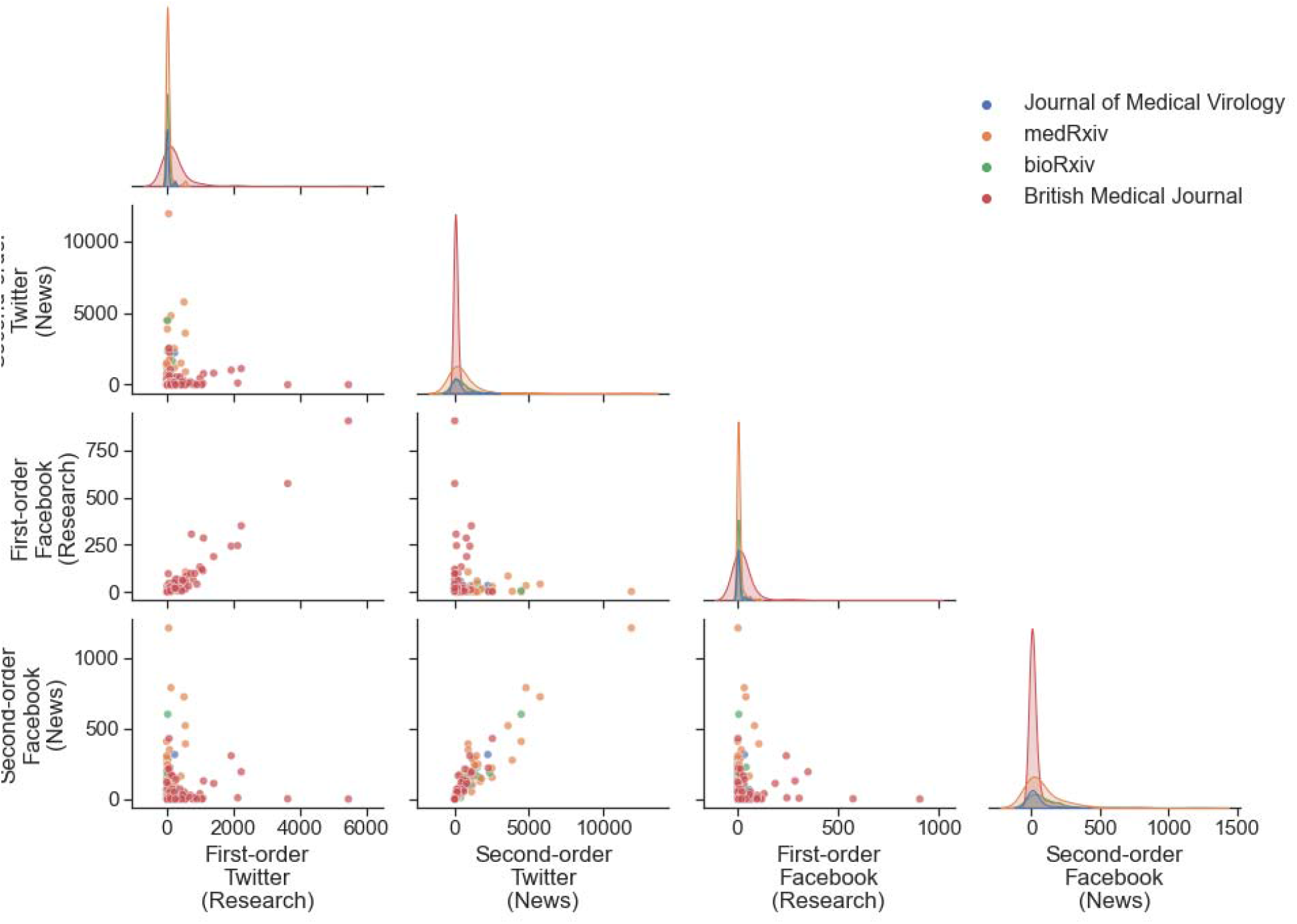
Comparison of first and second-order citations of research on Twitter and Facebook. Each data point represents a research article, with colour indicating publication venue.

### Most shared research

Table 4 lists the most popular research articles with the highest number of second-order citations on Twitter and Facebook in comparison to their first-order tweets and Facebook posts. Out of these 13 articles, 11 were preprints when they were mentioned in the news article and only 2 were already published in journals. However, 10 of the 11 preprints were later published in peer-reviewed journals, including well-established journals such as *Science, Nature, Cell*, and the *New England Journal of Medicine*. In line with the correlation patterns described above, only one of the research articles that was highly shared through second-order citations was also highly shared through first-order citations. Articles described findings with obvious practical value, such as assessments of the effectiveness of treatments or prevention measures [1,2,9] or situations that may increase one’s risk of contracting the virus [7,12,13]. Other highly shared news covered controversial subject matter such as social restrictions and hydroxychloroquine [1,2]. We further discuss possible factors contributing to the popularity of these articles in the Discussion below.

**Table 4.** Research articles with the most second-order citations on Twitter and Facebook

| Article Title | First-order<br>(cites research) |  |  |  | Second-order<br>(cites news) |  |  |  |
| --- | --- | --- | --- | --- | --- | --- | --- | --- |
|  | Tweets |  | Facebook |  | Tweets |  | Facebook |  |
|  | Num | Rank | Num | Rank | Num | Rank | Num | Rank |
| [1] Differential Effects of Intervention Timing on COVID-19 Spread in the United States (*medRxiv*; eventually in *Science Advances*) | 64 | 125 | 2 | 179 | 11,902 | 1 | 1,214 | 1 |
| [2] Outcomes of hydroxychloroquine usage in United States veterans hospitalized with Covid-19 (*medRxiv*; eventually in *Med*) | 528 | 23 | 41 | 32 | 5,754 | 2 | 726 | 3 |
| [3] Functional SARS-CoV-2-specific immune memory persists after mild COVID-19 (*bioRxiv*; eventually in *Cell*) | 136 | 69 | 33 | 37 | 4,796 | 3 | 790 | 2 |
| [4] Immunological memory to SARS-CoV-2 assessed for up to eight months after infection (*medRxiv*; eventually in *Science*) | 36 | 163 | 6 | 115 | 4,477 | 4 | 602 | 4 |
| [5] Sequencing identifies multiple early introductions of SARS-CoV-2 to the New York City Region (*medRxiv*; eventually in *Genome Research*) | 4 | 288 | 0 | 0 | 4,475 | 5 | 409 | 7 |
|  | Tweets |  | Facebook |  | Tweets |  | Facebook |  |
|  | Num | Rank | Num | Rank | Num | Rank | Num | Rank |
| [6] Coast-to-coast spread of SARS-CoV-2 in the United States revealed by genomic epidemiology<br>(medRxiv; eventually in *Cell*) | 28 | 181 | 3 | 157 | 3,876 | 6 | 276 | 16 |
| [7] Viable SARS-CoV-2 in the air of a hospital room with COVID-19 patients<br>(medRxiv; eventually in *International Journal of Infectious Diseases*) | 582 | 19 | 84 | 17 | 3,587 | 7 | 521 | 5 |
| [8] Covid-19: Local health teams trace eight times more contacts than national service<br>(*British Medical Journal*) | 77 | 109 | 2 | 179 | 2,534 | 8 | 430 | 6 |
| [9] Effect of Convalescent Plasma on Mortality among Hospitalized Patients with COVID-19: Initial Three-Month Experience<br>(medRxiv; not published elsewhere) | 247 | 43 | 29 | 43 | 2,520 | 9 | 153 | 40 |
| [10] Neutralising antibodies in Spike mediated SARS-CoV-2 adaptation<br>(medRxiv; eventually in *Nature*) | 54 | 139 | 4 | 141 | 2,498 | 10 | 220 | 23 |
| [11] The neuroinvasive potential of SARS-CoV2 may play a role in the respiratory failure of COVID-19 patients<br>(*Journal of Medical Virology*) | 255 | 40 | 34 | 35 | 2,228 | 13 | 316 | 10 |
| [12] Exposure to air pollution and COVID-19 mortality in the United States: A nationwide cross-sectional study<br>(medRxiv; eventually in *Science Advances*) | 95 | 95 | 19 | 51 | 910 | 39 | 349 | 9 |
| [13] Aerosol and surface stability of HCoV-19 (SARS-CoV-2) compared to SARS-CoV-1<br>(medRxiv; eventually in *New England Journal of Medicine*) | 609 | 18 | 106 | 12 | 896 | 40 | 392 | 8 |

### Overlaps in audiences

As shown in Figure 3, the overlap between the social media accounts that shared first- and second-order citations was very small on both Twitter (14.0% of 27,771 who shared research and 6.4% of 60,296 accounts that shared news stories) and Facebook (22.6% of 3,976 that shared research and 11.0% of 8,191 spaces that shared news stories).

**Figure 3.**
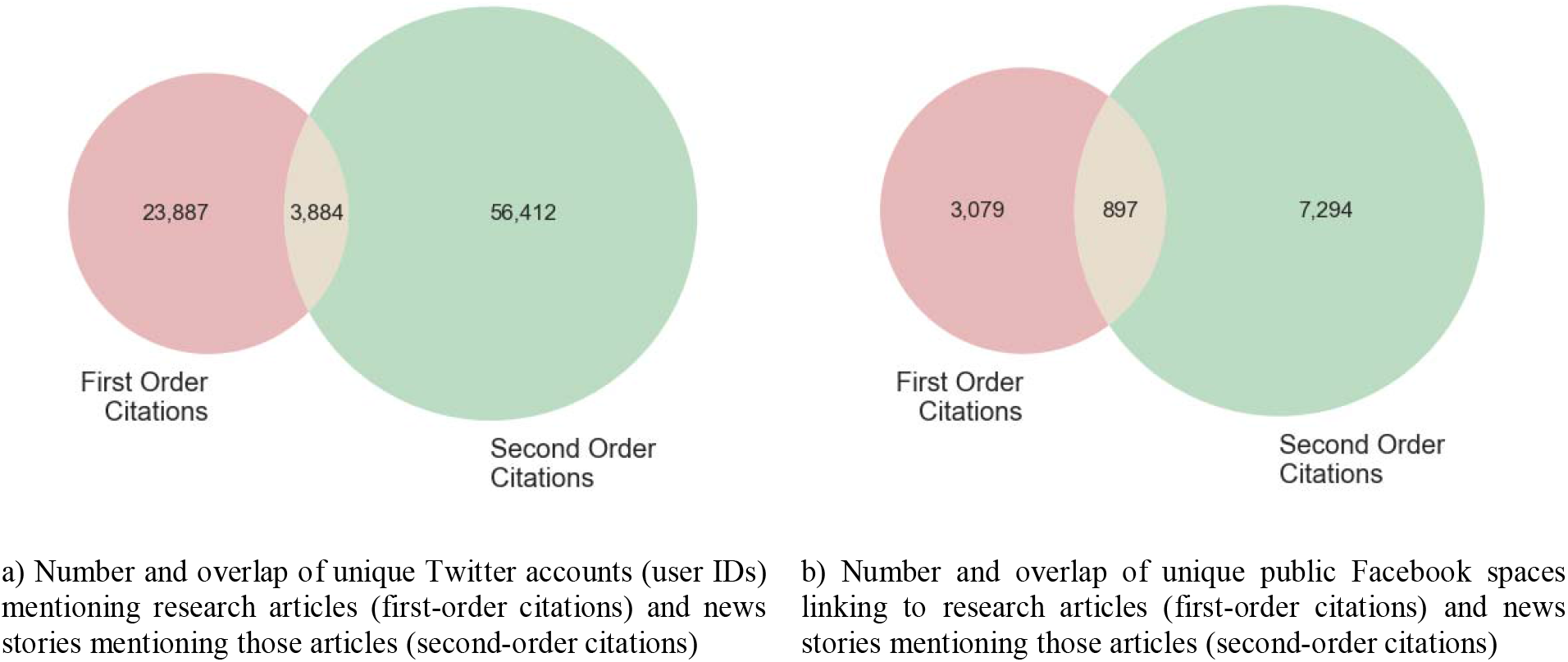
Number of unique accounts (Twitter) and public spaces (Facebook) linking to research articles (first-order citations) and news stories that mention those articles (second-order citations).

Table 5 lists the top 10 accounts with the most links to research (both first- and second-order citations) together with the top 10 accounts receiving the most engagement (likes and retweets). It includes highly visible scientists (e.g., Eric Topol, Eric Feigl-Ding), well-known journalists (e.g., Apoorva Mandavilli, George Monbiot) as well as political celebrity figures (i.e., Barack Obama, Hillary Clinton, Joe Biden). Results highlight drastically different levels of engagement that some social media accounts received. For example, we can see that Barack Obama’s single second-order citation (linking to an article in the *New York Times*) received 106,785 likes, which is more than all the likes received on tweets from any other account. For comparison, the account of the *British Medical Journal* (The BMJ) made 379 first-order citations that collectively received 41,068 likes (making it the fifth most liked account).

**Table 5.** Twitter accounts sharing the most links to research (first and second-order citations) and receiving the most user engagement (likes and retweets)

| Username | Name | Researcher* | Citation type | Tweets |  | Likes |  | Retweets |  |
| --- | --- | --- | --- | --- | --- | --- | --- | --- | --- |
|  |  |  |  | Rank | Num | Rank | Num | Rank | Num |
| bmj_latest | The BMJ | No | first | 1 | 379 | 5 | 41,068 | 1 | 27,263 |
| AndersJonita | Jonita Anders | No | first | 2 | 239 | 1,627 | 87 | 2,364 | 21 |
| uhiiman | 【自動化した】うひ<br>ーまん | No | second | 3 | 236 | 37,069 | 0 | 21,636 | 0 |
| Thomas_Wilckens | Dr. Thomas Wilckens | Yes | both | 4 | 226 | 512 | 406 | 403 | 211 |
| BangoBilly | //// Billy Bango \\\ | No | both | 5 | 197 | 339 | 664 | 286 | 312 |
| outbreaksci | Outbreak Science | No | first | 6 | 194 | 7,101 | 11 | 3,950 | 10 |
| BendallJane | jane bendall | No | both | 7 | 179 | 1,378 | 111 | 1,171 | 58 |
| Artaudculation | Leipzigconnection | No | both | 8 | 173 | 1,369 | 112 | 1,317 | 49 |
| pash22 | Ash Paul | Yes | both | 9 | 134 | 476 | 441 | 298 | 299 |
| tonto_1964 | Tony 'Gilets Jaunes' | No | first | 10 | 132 | 1,403 | 108 | 1,171 | 58 |
| EricTopol | Eric Topol | Yes | both | 37 | 64 | 10 | 22,953 | 8 | 10,914 |
| apoorva_nyc | Apoorva Mandavilli | Yes | both | 41 | 63 | 3 | 51,607 | 4 | 20,876 |
| trishgreenhalgh | Trisha Greenhalgh □<br>#CovidIsAirborne | Yes | both | 66 | 53 | 15 | 17,808 | 9 | 9,588 |
| DrEricDing | Eric Feigl-Ding | Yes | both | 272 | 26 | 4 | 46,702 | 5 | 15,736 |
| Karl_Lauterbach | Karl Lauterbach | No | both | 491 | 17 | 9 | 23,297 | 23 | 4,435 |
| GeorgeMonbiot | George Monbiot | No | second | 598 | 15 | 7 | 30,507 | 6 | 15,706 |
| carolecadwalla | Carole Cadwalladr | No | both | 2,233 | 6 | 6 | 32,878 | 7 | 15,229 |
| HillaryClinton | Hillary Clinton | No | second | 9,344 | 2 | 2 | 102,570 | 3 | 23,407 |
| neal_katyal | Neal Katyal | No | second | 9,344 | 2 | 11 | 22,183 | 10 | 8,808 |
| BarackObama | Barack Obama | No | second | 19,768 | 1 | 1 | 106,785 | 2 | 25,146 |
| JoeBiden | Joe Biden | No | second | 19,768 | 1 | 8 | 25,617 | 13 | 8,211 |

\*Included in the dataset published by Mongeon, Bowman, and Costas (2022)

Finally, we compared the Twitter accounts that had shared research and news with Twitter accounts of known researchers identified in the dataset published by Mongeon et al. (2022). We found small overlaps between the 423,920 Twitter accounts identified as belonging to a researcher and the accounts that made first- and second-order citations in our dataset, which can be explained by the focus on prioritizing precision over recall of Mongeon et al. (2023)’s approach. However, the share of researcher among accounts with first-order citations (14.0%; n=3,899) is more than twice as high as that among accounts with second-order citations (6.4%; n=3,830). As few as 718 accounts of the Mongeon et al. (2022) datasets were associated with both first- and second order citations in our dataset.

## Discussion

Numerous studies have found that hyperlinks to research articles shared on social media tend to circulate within closed communities of academic insiders rather than among wider public communities (Alperin et al., 2019; Carlson & Harris, 2020). The results of this study suggest that there may be other overlooked social media audiences that engage with research indirectly by sharing news coverage, or “second-order citations,” of research (Priem & Costello, 2010). Previous studies have suggested that these other audiences may be larger and more representative of society than audiences sharing research directly (Fleerackers, Riedlinger, et al., 2022; Lemke et al., 2021). Yet, to our knowledge, this is the first empirical study to document the relative sizes of these two audiences and to explore the degree to which they overlap with one another. The study also introduces a replicable method for identifying and analyzing the reach of second-order citations to research across two social media platforms, paving the way for more altmetrics research to consider indirect engagement with research.

Our findings suggest, albeit from a single case study, that expanding altmetric efforts to consider second-order citations is likely to be a productive avenue for identifying and understanding the circulation and impact of research beyond academic audiences. Our approach complements, and could be powerfully used in conjunction with, previous efforts to expand altmetrics from its origins of simply counting social media engagement with hyperlinks to research articles. In particular, we believe that an examination of second-order citations in conjunction with classification of social media accounts (Costas et al., 2020; Díaz-Faes et al., 2019; Fleerackers, Riedlinger, et al., 2022; Ke et al., 2017) and examination of social media network characteristics (Alperin et al., 2019; Costas et al., 2021; Robinson-Garcia et al., 2018) has the potential to yield greater insights about the public impact of research than what has been possible through the traditional focus on first-order citations.

This promise holds even as we acknowledge that the findings presented here cannot be generalized beyond the specific conditions of this case study of early COVID-19 research communication. Even with this caveat, the differences found between first- and second-order citations, in particular the lower share of researchers among those tweeting news stories rather than original articles, suggest that a broader segment of society may be engaging with research than what is commonly captured through first-order citations. In addition, we found that second-order citations also received far more social media engagement. Notably, this reach and engagement advantage was visible even when looking at just a small subset of available news coverage (i.e., stories published by only five media outlets). These findings suggest that online attention to second-order citations would likely be even larger when considering all news stories mentioning research. In other words, our findings indicate that second-order citations can reveal audiences of research that are significantly larger and have limited overlap with audiences identified through first-order citations, which was one of the initial goals of altmetrics (Priem et al., 2010).

An in-depth discussion of the causes behind the outsized engagement and reach of second-order citations is beyond the scope of this paper. However, potential factors to consider include journalists’ ability to make research knowledge more understandable and relevant to nonacademic audiences (Elliott, 2022; Gesualdo et al., 2020) and the public’s reliance on news coverage as a key source of science information (Funk et al., 2017)—particularly during the pandemic (Newman et al., 2020). Research into journalistic norms and routines may also help to explain the low correlations and lack of overlap between attention to first- and second-order citations. Specifically, journalistic notions of ‘relevance’ tend to focus on the usefulness or impact of research findings for society, while scholars tend to assess relevance based on whether research is novel or significant from an academic perspective (Elliott, 2022). Relatedly, journalists rely on well-established criteria when deciding what research to cover, selecting findings that have clear importance for their audience; that are surprising, novel, or controversial; or that are likely to spark curiosity and other emotions (Badenschier & Wormer, 2012; Rosen et al., 2016). In an age of metrics-driven news, journalists also work to maximize the online impact of their work, crafting “catchy” and clickable headlines and monitoring social media trends when selecting and framing stories (Moyo et al., 2019). Collectively, these journalistic practices may facilitate broader engagement with research findings on social media than is typically captured by altmetrics.

The imprint of journalistic selection criteria is visible in the list of highly shared research articles in our study (Table 4). Topics, such as treatment or prevention measures or high-risk situations to get infected have obvious practical values, which does not only align with traditional journalistic criteria for identifying ‘newsworthy’ science (Badenschier & Wormer, 2012), but also provides perfect fodder for the “news you can use” style of stories that appeared to be popular early on in the COVID-19 pandemic (Hermida et al., 2022). Other highly shared articles covered controversial topics, which align with journalistic selection criteria. For example, the preprint [1] that reported on the impacts of early non-pharmaceutical interventions on COVID-19 spread received 11,902 second-order citations on Twitter and so had huge social media impact through news media coverage, which might be due to the study’s societal relevance, emotional impact, and politicized media coverage.

The presence of Obama and other political celebrity figures, well-known journalists, and highly visible scientists in the top rankings (Table 5) aligns with previous research that has examined the role of individuals with celebrity status and other elites in the context of diffusing science content on social media (Gallagher et al., 2021; Joubert et al., 2023). Given what we know about the important role of influencer accounts in news curation (Bruns, 2018), it is perhaps unsurprising that these accounts generate engagement from sharing second-order citations, but it is also notable that 11 of the 21 accounts in the top rankings have shared both first- and second-order citations. While this analysis is not sufficient to make inferences about the role of identity and network position of those involved, it highlights how the identification of second-order citations and their circulation on social media platforms could be used to provide useful qualitative insights into the characteristics and identities of individuals who facilitate social media engagement with research.

Finally, the disconnects between what is popular as a first-order citation and what is popular as a second-order citation (described in Table 3 and shown in Figure 2), and the small overlaps in Twitter accounts and public Facebook spaces that are engaged with both types of citations (described in Figure 3), provide an additional rationale for examining second-order citations more closely. Both of these findings emphasize that a different set of individuals and mechanisms produce second-order citations, at least for the subset of COVID-19 research included in this case study.

## Conclusions

Although second-order citations were proposed over a decade ago (Priem & Costello, 2010), they remain unexplored by the altmetrics community. The findings presented in this article provide strong evidence for considering these indirect mentions of research as an important measure of impact beyond the academic community and so warrant further exploration. This is timely, given increasing recognition of the societal value of science communication and science journalism (Elliott, 2022; Gesualdo et al., 2020), not least because of their significant role in mobilizing knowledge during the COVID-19 pandemic (Joubert et al., 2023; Newman et al., 2020).

This paper also makes a significant contribution by pioneering a methodology for compiling and analyzing second-order citation data—something not currently available through existing altmetric data providers. Although recent changes at both Facebook and Twitter have likely jeopardized a direct replication of our method (Lawler, 2022; Weatherbed, 2023), the overall approach remains a valid one. While further work is required to test, extend, and evolve these methods, the findings described herein give us reason to believe that it will be worthwhile to do so.

## Appendix

Query from Chen et al. (2020) used to identify COVID-19 research in sample:

(((″BMJ (Clinical research ed.)″[Journal] OR (″Journal of medical virology″[Journal])) OR (″medRxiv : the preprint server for health sciences″[Journal])) OR (″bioRxiv : the preprint server for biology″[Journal])) AND ((″COVID-19″ OR ″COVID-19″[MeSH Terms] OR ″COVID-19 Vaccines″ OR ″COVID-19 Vaccines″[MeSH Terms] OR ″COVID-19 serotherapy″ OR ″COVID-19 serotherapy″[Supplementary Concept] OR ″COVID-19 Nucleic Acid Testing″ OR ″covid-19 nucleic acid testing″[MeSH Terms] OR ″COVID-19 Serological Testing″ OR ″covid-19 serological testing″[MeSH Terms] OR ″COVID-19 Testing″ OR ″covid-19 testing″[MeSH Terms] OR ″SARS-CoV-2″ OR ″sars-cov-2″[MeSH Terms] OR ″Severe Acute Respiratory Syndrome Coronavirus 2″ OR ″NCOV″ OR ″2019 NCOV″ OR ((″coronavirus″[MeSH Terms] OR ″coronavirus″ OR ″COV″) AND 2019/11/01[PDAT] : 3000/12/31[PDAT])))

## Acknowledgements

The initial idea for this study was jointly conceived by JPA, SH, and Vincent Larivière at the University of Montreal in the summer of 2015, for which we remain grateful nearly eight years later. We would also like to acknowledge Altmetric.com for access to the news media mentions data. Facebook data was provided by Crowdtangle through the API, and with the assistance of Jane Tan at the Digital Media Research Centre, Queensland University of Technology. Finally, we would like to thank the Health Communication Project team at the Scholarly Communication Lab for their input and support in the early stages of the research

## Funding

This work was supported by the Social Sciences and Humanities Research Council of Canada [435-2020-0401].

## Notes

https://zenodo.org/record/7803153

https://doi.org/10.7910/DVN/OEKB01

### Competing Interest Statement

The authors have declared no competing interest.

